# A characterization of the neural representation of confidence during probabilistic learning

**DOI:** 10.1101/2022.07.29.502028

**Authors:** Tiffany Bounmy, Evelyn Eger, Florent Meyniel

## Abstract

Learning in a stochastic and changing environment is a difficult task. Models of learning typically postulate that observations that deviate from the learned predictions are surprising and used to update those predictions. Bayesian accounts further posit the existence of a confidence-weighting mechanism: learning should be modulated by the confidence level that accompanies those predictions. However, the neural bases of this confidence are much less known than the ones of surprise. Here, we used a dynamic probability learning task and high-field MRI to identify putative cortical regions involved in the representation of confidence about predictions during human learning. We devised a stringent test based on the conjunction of four criteria. We localized several regions in parietal and frontal cortices whose activity is sensitive to the confidence of an ideal observer, specifically so with respect to potential confounds (surprise and predictability), and in a way that is invariant to which item is predicted. We also tested for functionality in two ways. First, we localized regions whose activity patterns at the subject level showed an effect of both confidence and surprise in qualitative agreement with the confidence-weighting principle. Second, we found neural representations of ideal confidence that also accounted for subjective confidence. Taken together, those results identify a set of cortical regions potentially implicated in the confidence-weighting of learning.

## Introduction

Many aspects of our environment are inherently stochastic, partly unknown, or so complex that we often cannot predict with certainty what will come next. Daily examples include the weather forecast, the duration of your commute and people’s behavior. Certainty is out of reach, but it often remains possible to characterize our environment by means of statistics, which is useful to guide our behavior. For instance, based on previous commute times, you can estimate the probability of a significant delay and plan accordingly when you really need to be on time.

The ability to extract statistics from previous observations is known as statistical learning, and it is known to be accurate in humans and other animals in a range of processes, from perception (Gallistel et al., 2014; Summerfield & de Lange, 2014; Saffran et al., 1996; Dehaene et al., 2015; Sherman et al., 2020), sensory-motor control (Wolpert et al., 1995; Berniker & Kording, 2008), to reinforcement learning (Sutton & Barto, 1998). Statistical learning remains accurate even in dynamic environments, in which statistics change over time (Iglesias et al., 2013; Peterson & Beach, 1967; Soltani & Izquierdo, 2019; Fusi et al., 2007).

In humans, learning is accompanied by a sense of confidence that characterizes the extent to which learners think that their learned estimates are accurate (Meyniel, Schlunegger, et al., 2015). Probabilistic inference (also referred to as Bayesian inference) provides an optimal solution to learn under uncertainty (Jaynes, 2003; Bland & Schaefer, 2012; Mathys, 2011). In the framework of probabilistic inference, the confidence about a learned estimate can be quantified by its precision (inverse variance), which accounts for several aspects of human confidence during statistical learning (Meyniel, Schlunegger, et al., 2015; Heilbron & Meyniel, 2019; Boldt et al., 2019; Nassar et al., 2010). Note that other aspects and formalizations of confidence exist in other tasks and contexts (see Discussion) and that here, confidence refers to the precision of a learned estimate.

Confidence seems to play a critical role when learning in dynamic environments (Nassar et al., 2012; Meyniel, Sigman, et al., 2015; Doya, 2002; Yu & Dayan, 2005; Vinckier et al., 2016; Courville et al., 2006). Previous studies showed that in such environments, humans (Behrens et al., 2007; Daw et al., 2006; Iglesias et al., 2013; Payzan-LeNestour et al., 2013) and other animals (Soltani & Izquierdo, 2019) dynamically adjust their learning. In probabilistic inference, the current learned estimate is updated all the more that new observations are surprising, and for a given level of surprise, this update is larger when confidence is lower. Several studies in humans, more indirectly in other animals, indicate that confidence is involved in the regulation of learning (Nassar et al., 2010; Grossman et al., 2022; Payzan-LeNestour et al., 2013; Frömer et al., 2021). Unraveling the neural representation of confidence would constitute a significant step to understand the neurobiological underpinning of learning under uncertainty.

However, a number of methodological challenges make the study of the neural representation of confidence difficult (Walker et al., 2022). First, confidence is not a property of some stimuli or the environment, but of an observer’s inference about their characteristics (like their statistics). The experimenter can ask participants to report their confidence and study its neural representation, but this approach is more likely to identify brain regions involved in the report of confidence than in its estimation and use in the learning process. An alternative is to not disrupt the learning process with reports, and instead rely on an (ideal observer) model of the learning process that quantifies confidence based on the sequence of stimuli. Here, we combine both approaches and use sequences of stimuli that are interrupted by reports only occasionally, and we use an ideal observer model to analyze the neural representation of confidence during stimulus presentation (between reports).

Another methodological challenge is that confidence is often confounded with other variables. For instance, surprise typically reduces confidence, and more predictable sequences (when the probability of the next item is more extreme) increase confidence. Here, we leveraged the use of an ideal observer model to quantify those confounding factors and test for the specificity of the neural representation of confidence with respect to those confounding variables.

A third methodological challenge is that the latent variables of the learning process can also be confounded with simple features of stimuli. For instance, if stimuli are generated based on a Gaussian distribution whose mean changes occasionally (Nassar et al., 2010; McGuire et al., 2014), then a stimulus that greatly differs from the previous one will be very surprising. Here, we simplified the stimulus space by using binary sequences. When there are only two possible items in a sequence, a stimulus can only repeat the previous one or change, and yet be associated with very different surprise and confidence levels depending on the statistical context.

The goal of the current study is to characterize the neural representation of confidence with four criteria often used in neuroscience (Walker et al., 2022). The activity of a candidate region representing confidence about a prediction should be sensitive to confidence, specifically so with respect to other variables, and in a way that is invariant to other features of the prediction (such as the identity of the predicted stimulus) that should in principle not impact confidence. The fourth criterion is functionality. As mentioned earlier, confidence is thought to regulate learning, with larger updates when surprise is larger and confidence lower. If true, then we expect that activity in some regions follows this pattern, and thus that some representations of confidence and surprise overlap anatomically (Meyniel & Dehaene, 2017; Meyniel, 2020). Another aspect of functionality is that the neural representation of (ideal) confidence during learning could inform the confidence reported by participants at the moment of questions, which we also tested.

We used fMRI to measure brain activity because it provides a whole brain coverage, which precludes in particular the need to rely on a priori regions of interest. We used 7T fMRI rather than a lower and more commonly used field (typically 3T), because it affords a higher signal to noise ratio, and thus a more precise anatomical characterization of neural representations, the possibility to perform subject-level analysis (which is critical for overlap analyses) and enough power to analyze even a small number of trials (like questions in our task).

## Results

### Bayesian inference accounts for subjects’ probability estimates and confidence levels

Twenty-six subjects performed a probability-learning task in an MRI scanner. Long sequences of visual stimuli, which were two Gabor patches oriented counterclockwise and clockwise (denoted A and B here for brevity) were presented to the subjects. These sequences were generated based on a Bernoulli process, in which the item probability p(A) (and thus p(B)=1-p(A)) was constant between change points that occurred at random, unsignaled moments. Subjects were informed of the task structure and were asked to covertly estimate p(A). We inserted a few question trials in the sequence for participants to report their prediction about the next item in the form of a probability estimate, and their confidence about this estimate (see Figure 1). We compared those reports to those of an ideal observer model that uses probabilistic inference and previous observations in the sequence to estimate (optimally) a distribution of the item probability p(A), from which it reads the prediction as the mean and confidence as the log precision (see Methods). We first computed the correlation between participants’ responses and the ideal observer (for both probability estimates and confidence ratings), then analyzed the group-level correlations. Results showed that participants’ probability estimates linearly correlated with the ideal observer estimates (ρ=0.79, SD=0.126, SEM=0.025, Cohen’s d=6.231, t_25_=31.8, P=9.99 x 10^-22^, see Figure 1 bottom left) and subjective confidence levels linearly correlated with the (ideal observer) confidence (ρ=0.16, SD=0.134, SEM=0.026, Cohen’s d=1.160, t_25_=5.9, P=3.6 x 10^-6^, see Figure 1, bottom right). Those results are in line with previous studies testing subjects on a similar task (Meyniel, Schlunegger, et al., 2015; Meyniel & Dehaene, 2017; Heilbron & Meyniel, 2019; Meyniel, 2020).

**Figure 1.**
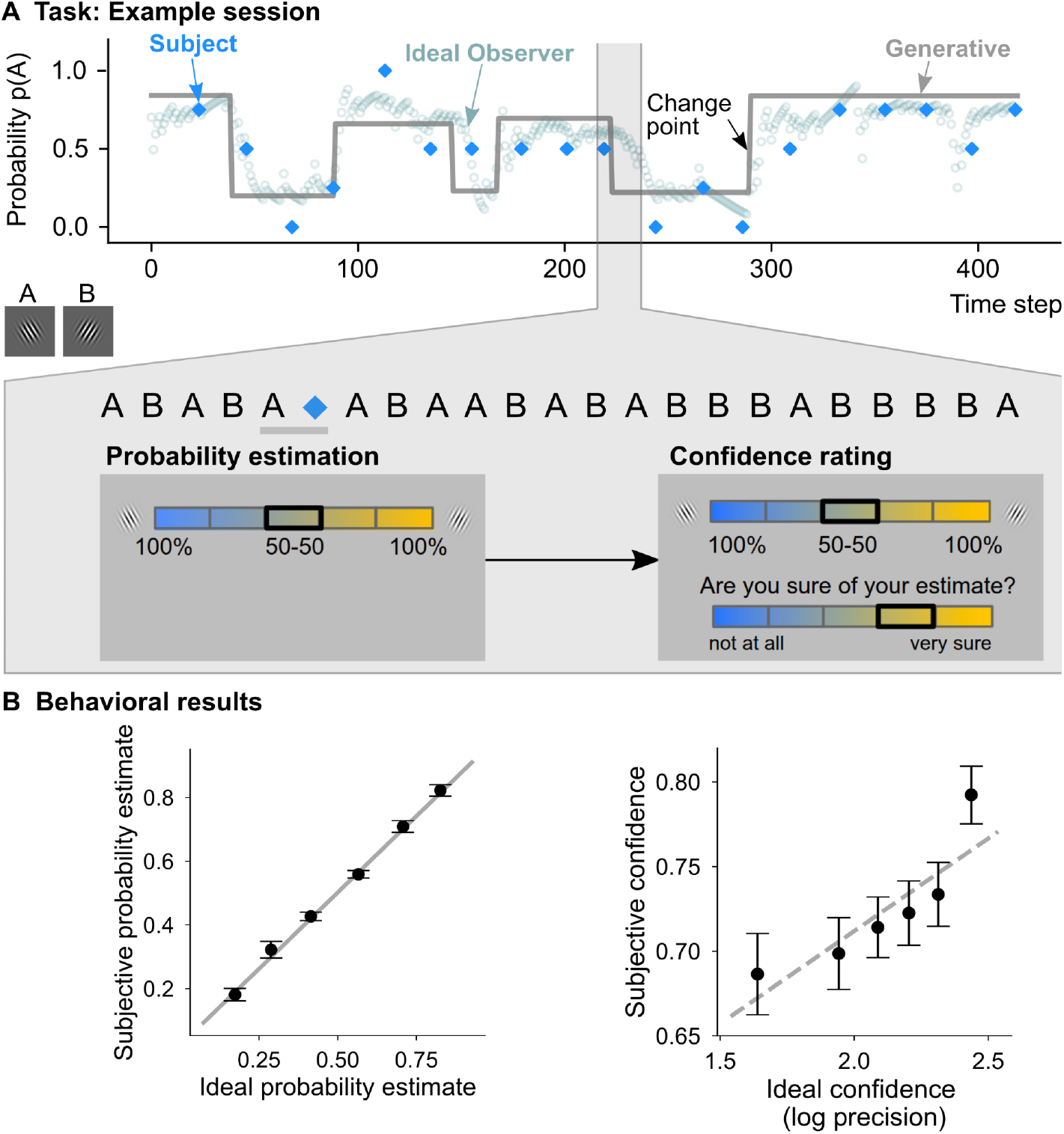
Bayesian inference accounts for subjective probability estimates and confidence in a dynamic probability-learning task. **A.** Example session of the probability-learning task. The sequence was composed of two Gabor patches (oriented counterclockwise and clockwise, and denoted A and B respectively), as illustrated by the snippet. In each session a sequence showed 420 stimuli (one stimulus every 1.3 s) drawn from a Bernoulli process in which the item probability p(A) changed abruptly and unpredictably. Occasionally (every 22 stimuli on average), the sequence was interrupted and participants reported their estimate of p(A) and their confidence about their estimate (the order of questions was counterbalanced across participants; 5 probability bins are shown in this example question screen but 3 bins were also used, see Methods). The gray line is the generative p(A), circles show the estimates of an ideal observer and the blue diamonds show the subject’s probability estimates. **B.** Reported probability estimates and confidence levels sorted in 6 bins of ideal estimates; dots and error bars indicate the mean and S.E.M. The plain line indicates the identity, the dashed line is a linear fit. Note that subjective confidence (lowest and highest response bins set to 0 and 1 arbitrarily) and ideal confidence (in log SD unit) are on different scales.

Subjective confidence, although less correlated than predictions to the ideal observer, showed the two following signatures of a probabilistic inference. Two variables are known to influence confidence in probabilistic inference (Meyniel, Schlunegger, et al., 2015; Nassar et al., 2010): unpredictability (how unsure the ideal observer is about the identity of the next item, this uncertainty increases as the estimated p(A) approaches 0.5) and surprise (the extent to which the last observation deviated from the estimated p(A), see Methods). In the ideal observer, there are strong negative correlations indeed between confidence and unpredictability (ρ=-0.31, SD=0.121, SEM=0.024, Cohen’s d=-2.546, t_25_=-13.0, p=1.3 x 10^-12^), and between confidence and surprise (ρ=-0.61, SD=0.086, SEM=0.017, Cohen’s d=-7.167, t_25_=-36.5, P=3.2 x 10^-23^). Qualitatively similar effects were found in subjects (correlation between subjective confidence and ideal estimate of unpredictability: ρ=-0.28, SD=0.143, SEM=0.028, Cohen’s d=-1.970, t_25_=-10.00, P=2.9 x 10^-10^; correlation between subjective confidence and ideal surprise: ρ=-0.11, SD=0.120, SEM=0.024, Cohen’s d=-0.950, t_25_=-4.8, P=5.6 x 10^-5^, see Figure 1—figure supplement 1). Those correlations highlight the quality of subjective confidence reports, but they also indicate that surprise and unpredictability are confounding factors in the analysis of confidence. The fMRI analysis below takes those confounding factors into account.

### Brain networks specifically sensitive to confidence and surprise

The fMRI analysis focused on the sequence of stimuli between questions, when subjects covertly estimate probabilities, which leaves brain activity unperturbed by the act of reporting an estimate. We regressed several variables (confidence, surprise, unpredictability, prediction) derived from the ideal observer model onto fMRI activity to explore the neural substrates of probabilistic inference, knowing that the ideal observer accounts for the subjects’ inference (see previous section). This analysis assumes that a region that represents confidence has activity levels that increase and decrease following confidence; this assumption is frequent in the fMRI literature (see Discussion for the implication of this assumption). Before modeling fMRI activity, we performed a parameter recovery analysis in order to test the extent to which our effects of interest (confidence and surprise) can be uniquely estimated with a general linear model (GLM_1_) when other confounds (predictability and the prediction itself) are included in the model. The simulation of 1000 fMRI-like experiments showed that we can recover the parameters even in the presence of strong autocorrelated noise (see Figure 2A).

**Figure 2.**
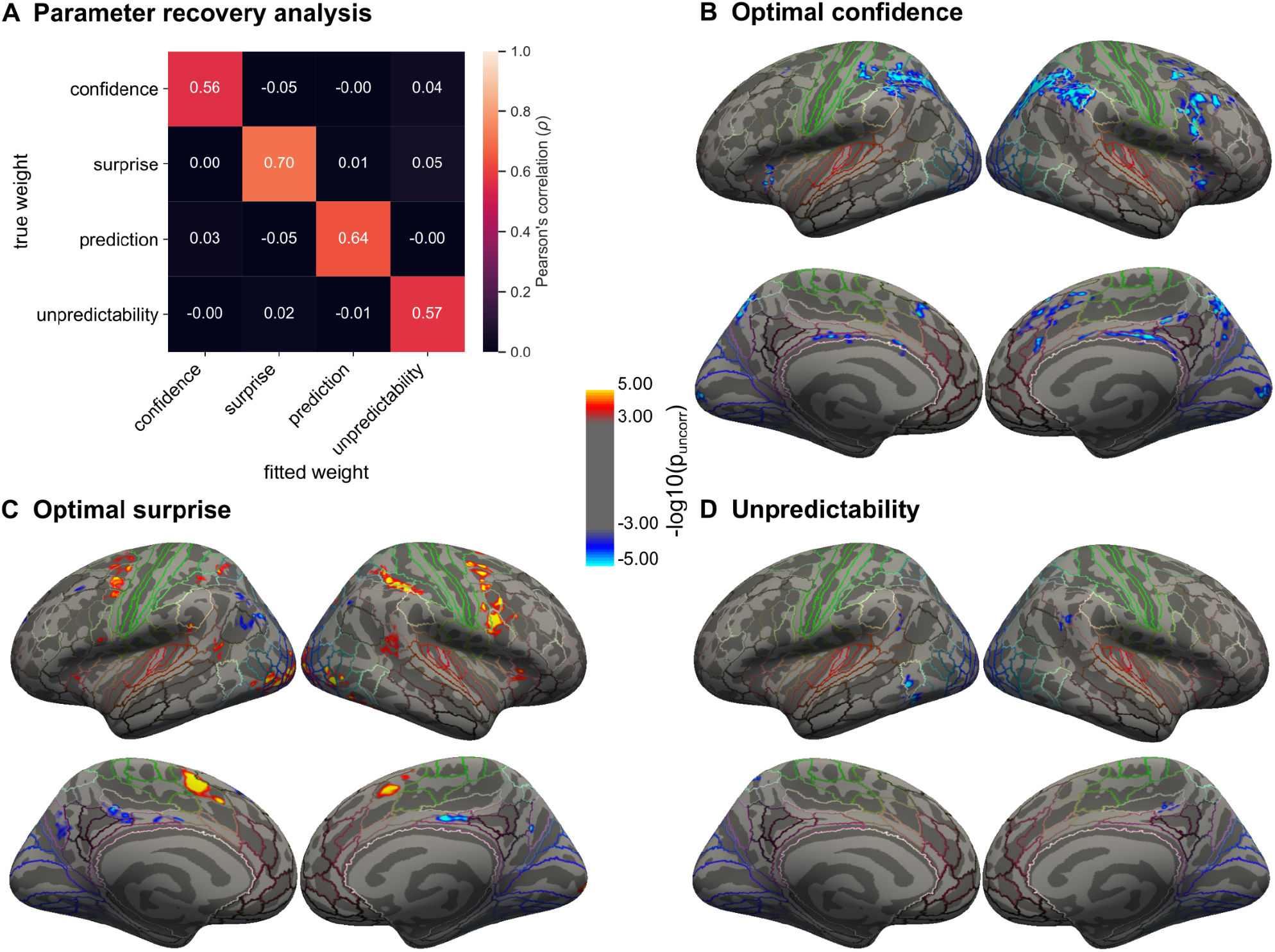
Activity in a fronto-parietal network is sensitive and specific to different latent variables of learning (confidence, surprise and unpredictability). **A.** Parameter recovery analysis. Pearson’s correlation between the true and fitted weights of the latent variables of learning (confidence, surprise, unpredictability and prediction) in GLM_1_.The simulated fMRI signal was corrupted with strong autocorrelated noise (AR1=0.9 with an average ratio of 0.10 between noise-free and noisy signals’ SD). **B-D.** Brain regions where activity is significantly correlated with confidence (**B**), surprise (**C**) and unpredictability (**D**). Significance maps (-log p-value) from the surface-based, group-level analysis (GLM_1_) were thresholded at a cluster-forming threshold of p<0.001 and corrected for multiple comparisons across vertices at the cluster level with p_FWE_<0.05. Anatomical landmarks correspond to the HCP-MMP1.0 atlas (Glasser et al., 2016).

With the actual fMRI data, the analysis (GLM_1_, see Methods) revealed the existence of multiple regions in fronto-parietal cortices whose activity correlates with confidence, surprise and unpredictability (see Figure 2B-D). More precisely, we located several clusters correlating negatively with confidence (Figure 2B, Table 2), notably in the superior parietal cortex that included the right intraparietal sulcus [*x*, *y, z:* (27, −61, 41)] and left intraparietal area 2 [*x, y z:* (−46, −45, 41)], the right inferior frontal cortex [*x, y, z:* (38, 10, 22)] and the primary visual cortex [*x, y, z:* right (14, −8, 1), left (−14, −92, 0.5)]. We also identified several clusters correlating positively with surprise (Figure 2C; Table 1), notably in the right intraparietal sulcus, the right frontal eye field, the right superior frontal gyrus and the occipital pole, and negatively in the inferior temporal gyrus. Note that GLM_1_ is suitable to assess the specificity of the neural correlates of confidence with respect to confounding variables like surprise and unpredictability because those variables are included in the same model. In particular, the effects of unpredictability were anatomically distinct from those of confidence: fMRI activity negatively correlated with unpredictability in much smaller portions of the cortex, in the right inferior parietal gyrus, the right marginal sulcus, the left lateral occipitotemporal gyrus, the left inferior and superior parietal gyri and the left middle temporal gyrus (Figure 2D). Significance maps, uncorrected for multiple comparisons and unthresholded (Figure 2—figure supplement 1) suggest that other regions may be involved, but with more limited statistical significance, like the ventral medial prefrontal and frontopolar cortices whose activity correlated positively with confidence. In addition, we verified that similar regions are identified for the effects of surprise and confidence in the two groups of participants (N=13 each) who answered the probability and confidence questions in a different order (see unthresholded significance maps Figure 2—figure supplement 2). This comparison serves as an internal replication of the results, and indicates that the neural correlates of confidence and surprise do not depend much on the specifics of the reporting procedure. Last, we also verified that the brain regions identified so far for confidence and surprise at the group-level analysis can be found at the individual level, for some participants at least (see example in Figure 2—figure supplement 3).

**Table 1.**
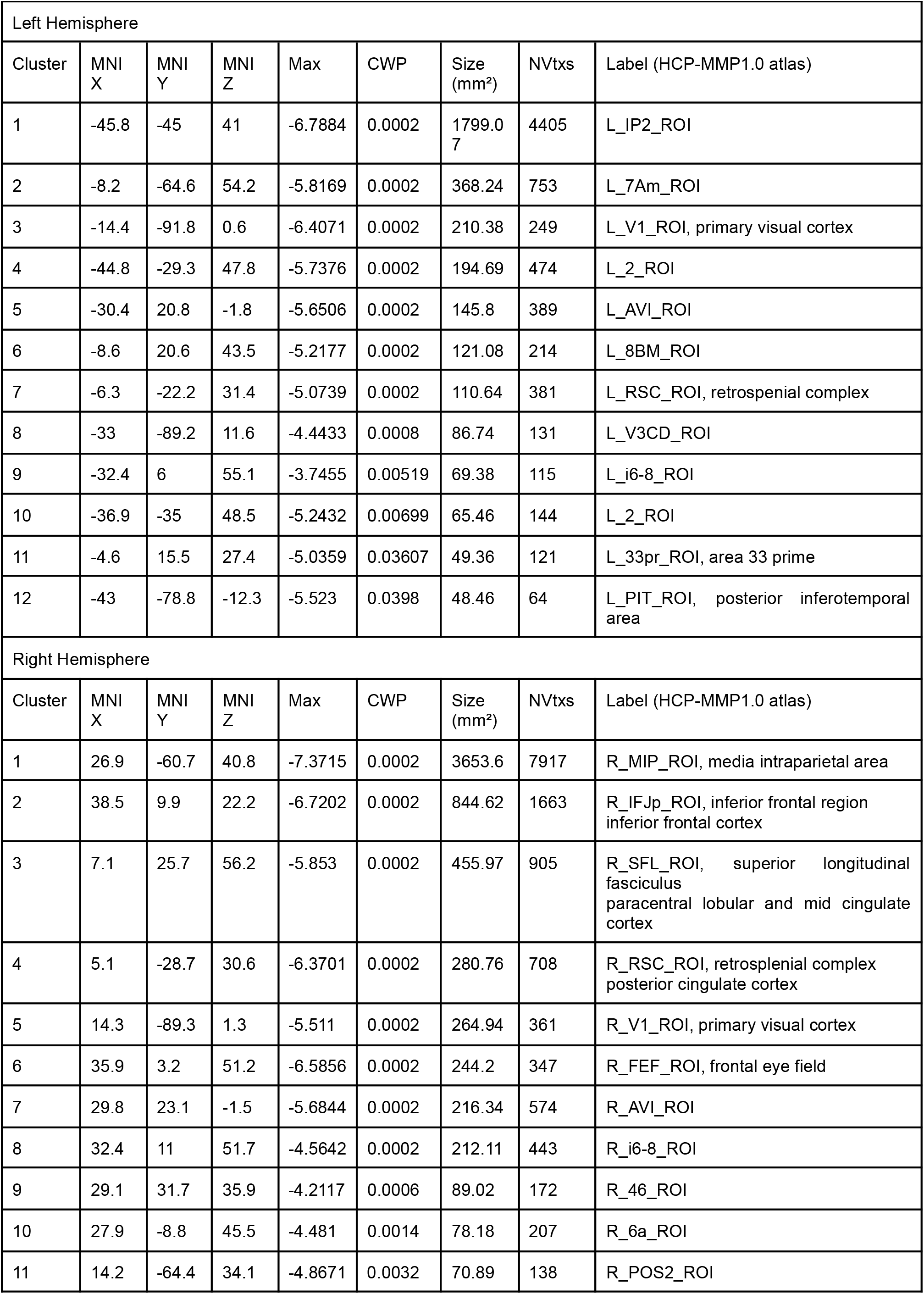
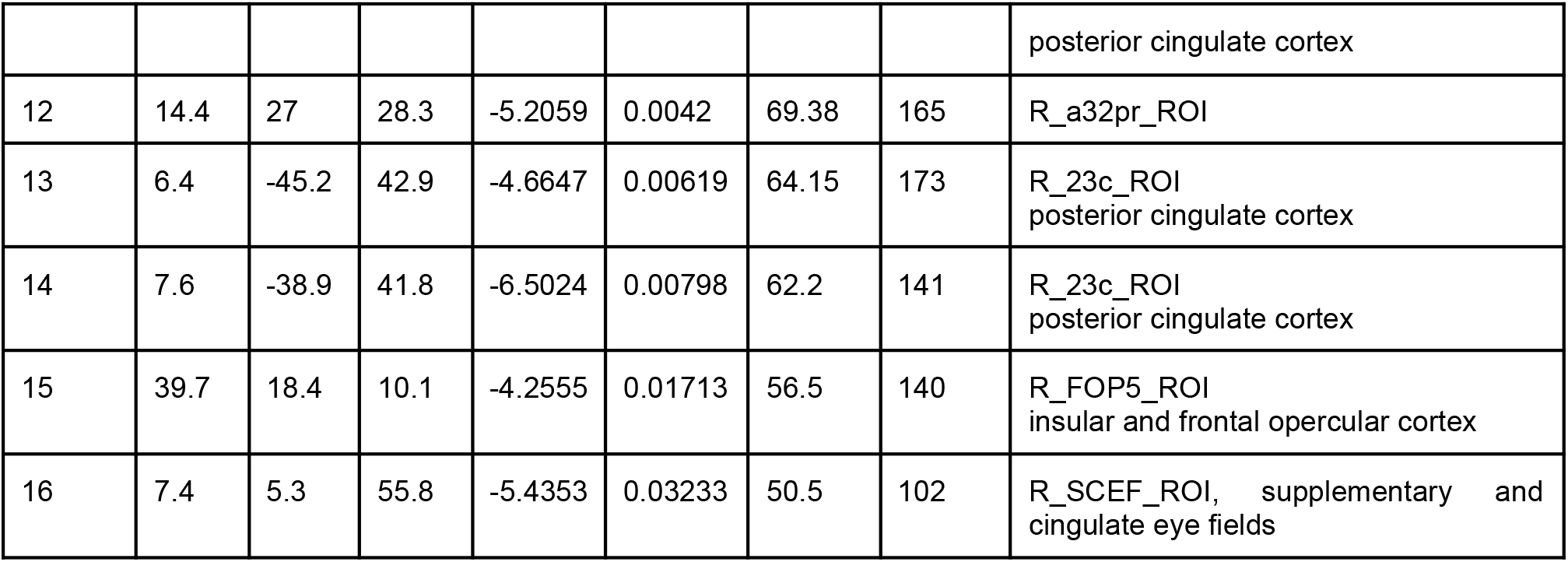
Cortical effects of confidence.

**Table 2.** Cortical effects of surprise.

| Left Hemisphere |  |  |  |  |  |  |  |  |
| --- | --- | --- | --- | --- | --- | --- | --- | --- |
| Cluster | MNI X | MNI Y | MNI Z | Max | CWP | Size (mm²) | NVtxs | Label (HCP-MMP1.0 atlas) |
| 1 | -28.9 | -91.1 | -5.6 | 7.047 | 0.00020 | 1628.19 | 2157 | L_V3_ROI |
| 2 | -43.9 | 0.4 | 46.5 | 5.7707 | 0.00020 | 734.29 | 1532 | L_55b_ROI |
| 3 | -6.5 | 1.0 | 61.7 | 6.4885 | 0.00020 | 427.31 | 825 | L_SCEF_ROI |
| 4 | -41.6 | -68.7 | 29.4 | -5.3645 | 0.00020 | 287.09 | 560 | L_PGs_ROI<br>(inferior parietal cortex) |
| 5 | -9.6 | -65.6 | 27.5 | -3.9612 | 0.00020 | 228.06 | 446 | L_POS2_ROI |
| 6 | -6.9 | 28.0 | 55.0 | -4.4231 | 0.00020 | 225.00 | 389 | L_SFL_ROI |
| 7 | -10.6 | -37.6 | 41.4 | -5.7393 | 0.00020 | 217.42 | 498 | L_23c_ROI |
| 8 | -40.1 | -64.3 | 44.9 | -4.351 | 0.00020 | 199.53 | 434 | L_PFm_ROI |
| 9 | -32.8 | -51.3 | 53.2 | 5.0564 | 0.00020 | 169.69 | 393 | L_LIPv_ROI, lateral intraparietal area |
| 10 | -46.7 | -48.1 | 12.7 | 4.5669 | 0.00020 | 114.47 | 248 | L_TPOJ1_ROI, temporo-parieto-occipito<br>junction |
| 11 | -31.3 | -40.4 | 42.5 | 4.4284 | 0.00020 | 104.49 | 302 | L_AIP_ROI, anterior intraparietal area |
| 12 | -41.7 | -66.7 | -14.9 | 5.7507 | 0.00260 | 78.21 | 115 | L_FFC_ROI, fusiform face complex |
| 13 | -4.0 | -4.1 | 34.3 | -5.1037 | 0.00778 | 64.59 | 175 | L_23d_ROI |
| 14 | -36.9 | 22.5 | 45.3 | -4.1085 | 0.01117 | 60.55 | 99 | L_8Av_ROI |
| 15 | -48.0 | -40.7 | 27.5 | 5.09 | 0.01990 | 54.39 | 137 | L_PF_ROI |
| 16 | -20.4 | 49.1 | 27.7 | -4.0626 | 0.02465 | 53.13 | 83 | L_9a_ROI |
| 17 | -33.2 | -77.7 | 39.2 | -3.7778 | 0.02524 | 52.92 | 82 | L_PGs_ROI |
| 18 | -49.3 | 4.7 | 10.5 | 4.2695 | 0.03273 | 50.90 | 116 | L_6r_ROI |
| 19 | -27.8 | 24.5 | 2.6 | 3.9097 | 0.03921 | 48.53 | 143 | L_AVI_ROI |
| 20 | -46.4 | -65.5 | -4.9 | 4.4998 | 0.04312 | 47.95 | 86 | L_FST_ROI |

| Right Hemisphere |  |  |  |  |  |  |  |  |
| --- | --- | --- | --- | --- | --- | --- | --- | --- |
| Cluster | MNI X | MNI Y | MNI Z | Max | CWP | Size (mm<sup>2</sup>) | NVtxs | Label (HCP-MMP1.0 atlas) |
| 1 | 41.7 | -76.5 | -2.9 | 6.2505 | 0.0002 | 1200.64 | 1568 | R_V4t_ROI |
| 2 | 37.2 | -40.5 | 35.3 | 6.6126 | 0.0002 | 846.61 | 2117 | R_AIP_ROI |
| 3 | 50.2 | 6.5 | 27.9 | 7.0213 | 0.0002 | 635.71 | 1312 | R_6v_ROI |
| 4 | 28.1 | -3.2 | 45.6 | 6.0902 | 0.0002 | 570.11 | 1062 | R_6a_ROI |
| 5 | 48 | -43.7 | 16.5 | 4.3468 | 0.0002 | 304.23 | 806 | R_STV_ROI |
| 6 | 44.5 | -63 | -10.8 | 5.5522 | 0.0002 | 253.44 | 406 | R_PH_ROI |
| 7 | 26.2 | -92.2 | -12.6 | 4.1146 | 0.0002 | 249.49 | 293 | R_V3_ROI |
| 8 | 16.5 | -98.9 | -3.4 | 8.3045 | 0.0002 | 173.81 | 229 | R_V1_ROI |
| 9 | 4.1 | -18.5 | 36.8 | -5.7099 | 0.0002 | 148.18 | 342 | R_23d_ROI |
| 10 | 8.4 | 7.7 | 53 | 5.5549 | 0.0002 | 132.59 | 266 | R_SCEF_ROI, supplementary and cingulate eye fields |
| 11 | 40.6 | -68 | 39.2 | -4.0854 | 0.0002 | 114.92 | 198 | R_PGs_ROI |
| 12 | 17.6 | -0.3 | 65.4 | 4.979 | 0.0004 | 97.52 | 207 | R_6ma_ROI |
| 13 | 31 | 27.3 | 6.7 | 4.7609 | 0.0014 | 80.95 | 221 | R_FOP5_ROI |
| 14 | 14.9 | -69.5 | 53.2 | 4.5225 | 0.0014 | 77.89 | 134 | R_7PL_ROI |
| 15 | 7.6 | -1 | 65.6 | 4.3585 | 0.002 | 74.51 | 137 | R_SFL_ROI |
| 16 | 26.7 | -68.4 | 30.1 | 4.4314 | 0.0024 | 73.31 | 94 | R_IPS1_ROI |
| 17 | 10.8 | 38.1 | 49.4 | -4.6485 | 0.00739 | 62.33 | 110 | R_8BL_ROI |
| 18 | 7.4 | -66.9 | 29.4 | -5.5523 | 0.03312 | 50.18 | 94 | R_POS2_ROI |

**Table 3.** Cortical effects of unpredictability.

| Left Hemisphere |  |  |  |  |  |  |  |  |
| --- | --- | --- | --- | --- | --- | --- | --- | --- |
| Cluster | MNI X | MNI Y | MNI Z | Max | CWP | Size (mm<sup>2</sup>) | NVtxs | Label (HCP-MMP1.0 atlas) |
| 1 | -46 | -54.1 | -9.3 | -6.0018 | 0.0002 | 123.64 | 194 | L_PH_ROI |
| 2 | -56.2 | -52.4 | -4 | -5.6255 | 0.0002 | 113.34 | 189 | L_PHT_ROI |
| 3 | -13.7 | -60.4 | 59.3 | -5.1744 | 0.01177 | 59.89 | 136 | L_7Am_ROI |
| 4 | -57.3 | -51.7 | 20.6 | -4.6458 | 0.0201 | 54.2 | 130 | L_PFm_ROI |
| Right Hemisphere |  |  |  |  |  |  |  |  |
| Cluster | MNI X | MNI Y | MNI Z | Max | CWP | Size (mm<sup>2</sup>) | NVtxs | Label (HCP-MMP1.0 atlas) |
| 1 | 53.3 | -47.4 | 27 | -4.7213 | 0.0002 | 113.63 | 251 | R_PFm_ROI |
| 2 | 10.7 | -38.6 | 40.9 | -4.5575 | 0.0004 | 96.21 | 234 | R_23c_ROI |

Altogether, those results show that fMRI activity is sensitive to the latent variables of learning, and also specific given that confidence, surprise and unpredictability were all included in GLM_1_ and several clusters associated with each effect are distinct.

### Neural correlates of confidence are invariant to the predicted item

We then tested whether the neural correlates of confidence are invariant to the predicted item, in other words, whether fMRI activity is similar for a given level of confidence, no matter whether this confidence is associated with the prediction of a Gabor patch oriented clockwise or counterclockwise. To test for this invariance, we estimated the effect (whole-brain) of confidence separately for stimuli when the predicted item in the sequence was oriented clockwise or counterclockwise, entered as two regressors in GLM_2_ (see Methods). We found similar regions for effects of confidence in both cases, in particular in the intra-parietal cortex and the inferior frontal gyrus identified previously (Figure 2), see Figure 3—figure supplement 1. In addition, we tested for invariance systematically in the clusters identified previously for confidence (Figure 2) by extracting their fMRI activity for stimuli sorted into different confidence bins, and further sorted into whether the current prediction favored a Gabor patch oriented clockwise or counterclockwise (GLM_3_). For the sake of simplicity, neighboring clusters within an hemisphere, and homologous clusters across the two hemispheres, were merged. The results (Figure 3) showed that most clusters, notably in the superior parietal cortex, the insular and fronto opercular cortex and the mid cingulate cortex, showed similar confidence effects irrespective of the predicted Gabor patch orientation. Those results also indicate that the negative correlation identified in Figure 2 corresponds to graded levels of fMRI activity for graded levels of confidence, rather than being driven by extreme values.

**Figure 3.**
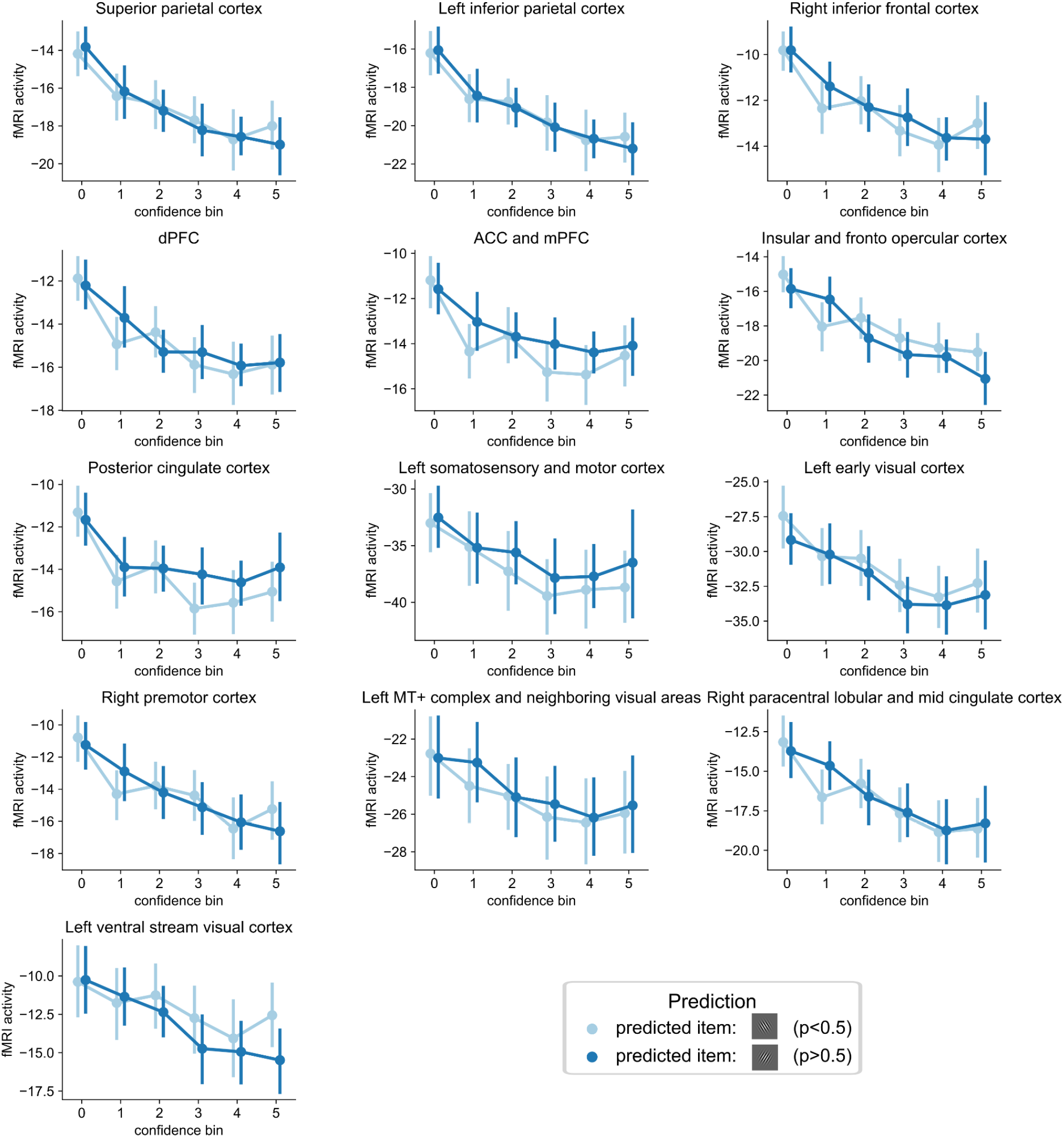
Invariance of the confidence effect with respect to the predicted item. Beta coefficients (first-level estimates) for different confidence bins. Relative fMRI activity in bins of confidence, sorted by the predicted item in the sequence (oriented counterclockwise (p(clockwise)<0.5) or clockwise (p(clockwise)>0.5)), obtained with GLM_3_. Dots and error bars indicate the mean and S.E.M. Regions are sorted (left-to-right on a given row and top-to-bottom across rows) following the strength of the confidence effect.

### Candidate regions for probability updates: overlap between surprise and confidence effects

In (optimal) probabilistic inference, the updating process of the current estimates takes into account both confidence and surprise. More surprising observations lead to updating the current estimate more, and for a given surprise level, this update is all the smaller that confidence about the prediction was higher (Jaynes, 2003; Gelman et al., 2013; McGuire et al., 2014; Meyniel & Dehaene, 2017). A brain region involved in this update may thus exhibit larger fMRI activity when confidence is lower and when surprise is larger. Looking for candidate regions for probability update by means of a correlation with the amount of update is insufficient because such a correlation can be driven by either confidence or surprise alone. A more powerful test is to look for the existence of an overlap of the brain regions that correlate positively with surprise and those that correlate negatively with confidence (Figure 2) which we detected at the group-level, notably in the intraparietal cortex and inferior frontal gyrus (see Figure 4A). Those regions could thus contribute to updating the current probability estimate.

**Figure 4.**
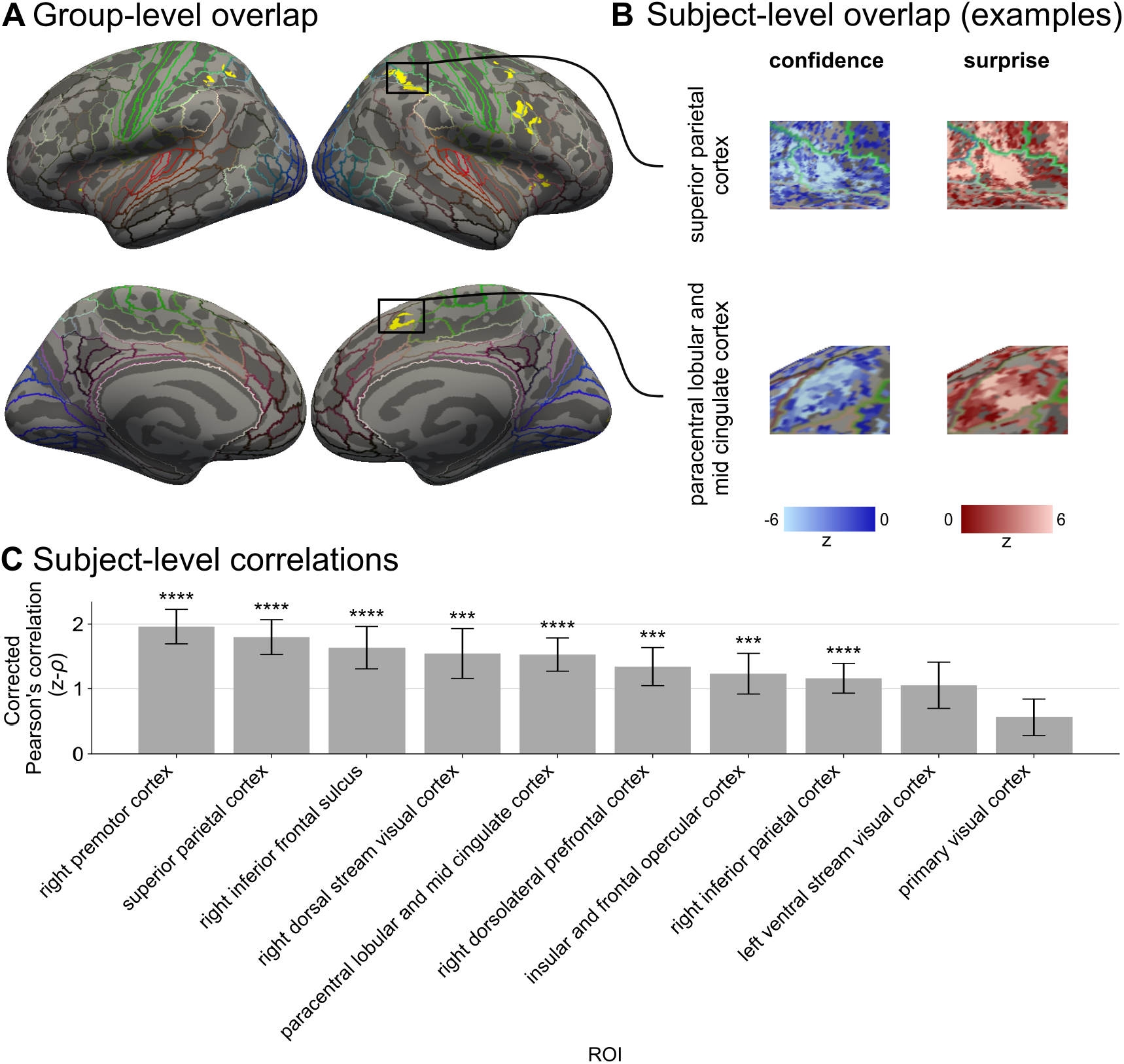
Candidate regions for the update of probability estimates. **A.** Group-level overlap of the effects of surprise and uncertainty (yellow), each significance map being thresholded at a cluster-forming threshold of p<0.001 and corrected for multiple comparisons across vertices at the cluster level with p_FWE_<0.05. **B.** Example of overlap between the effects of surprise and uncertainty in an example subject. **C.** Standardized correlation (z-ρ) estimated at the subject-level between the effect size of surprise and uncertainty in different ROIs; bars and error bars show the means and S.E.M. ****: p<0.0001; ***: p<0.001.

However, the anatomical overlap of those two effects at the group level may actually arise from different groups of subjects exhibiting an effect of confidence and surprise in the same region. To determine whether the overlap of surprise and confidence effects in those regions can also be found at the subject level, we measured for each subject the correlation between the regression coefficients of the effects of surprise and confidence (GLM_1_) across the vertices of each cluster showing a significant group-level overlap of those two effects. The underlying assumption is that if there is a subject-level overlap of those effects in a region, a vertex that exhibits a stronger (respectively, weaker) effect of one factor should also exhibit a stronger (respectively, weaker) effect of the other factor (L. Wang et al., 2015). To facilitate the interpretation of numerical values, we used uncertainty (the negative confidence) and surprise, since a region involved in the update should exhibit a positive correlation with both factors. To compare the observed correlation with the chance-level correlation, we standardized the result (using a null distribution where no relationship is expected between surprise and confidence on the one hand, and fMRI activity on the other, see Methods). The standardized correlation was positive (but not always significant) in all clusters showing a group-level overlap indicating the existence of an overlap at the subject level, which was particularly strong in the intraparietal cortex, the precentral-inferior frontal gyrus (Figure 4B). Examples of subject-level maps of those two effects exhibiting an overlap are shown in Figure 4C.

### Confidence networks are functional

So far, we have analyzed the effect of the confidence of an ideal observer on fMRI activity. If the brain regions identified with this analysis have a functional role, their activity may also relate to the confidence reported by participants at the moment of questions. More precisely, following the direction of the effect observed with the ideal observer, the neural activity in those regions before the onset of questions should be larger when subjects report being more uncertain about their estimate. To test for this effect in the confidence clusters identified with GLM_1_, we performed a finite-impulse response (FIR) analysis of fMRI activity in a time window comprising the onset of the question screen, sorting questions into those with low vs. high subjective confidence. This analysis takes into account the confidence of the ideal observer during the sequence of stimuli (GLM_4_, see Methods). At the group-level, a number of regions exhibited the expected difference in fMRI activity depending on the confidence reported by subjects already at the time of the question onset (see Figure 5). Given the delay between neural activity and fMRI signals, this result indicates that neural activity before the question onset in those regions is indicative of the confidence that the participant will report, which is in favor of a functional role of those regions’ clusters in the estimation or use of confidence.

**Figure 5.**
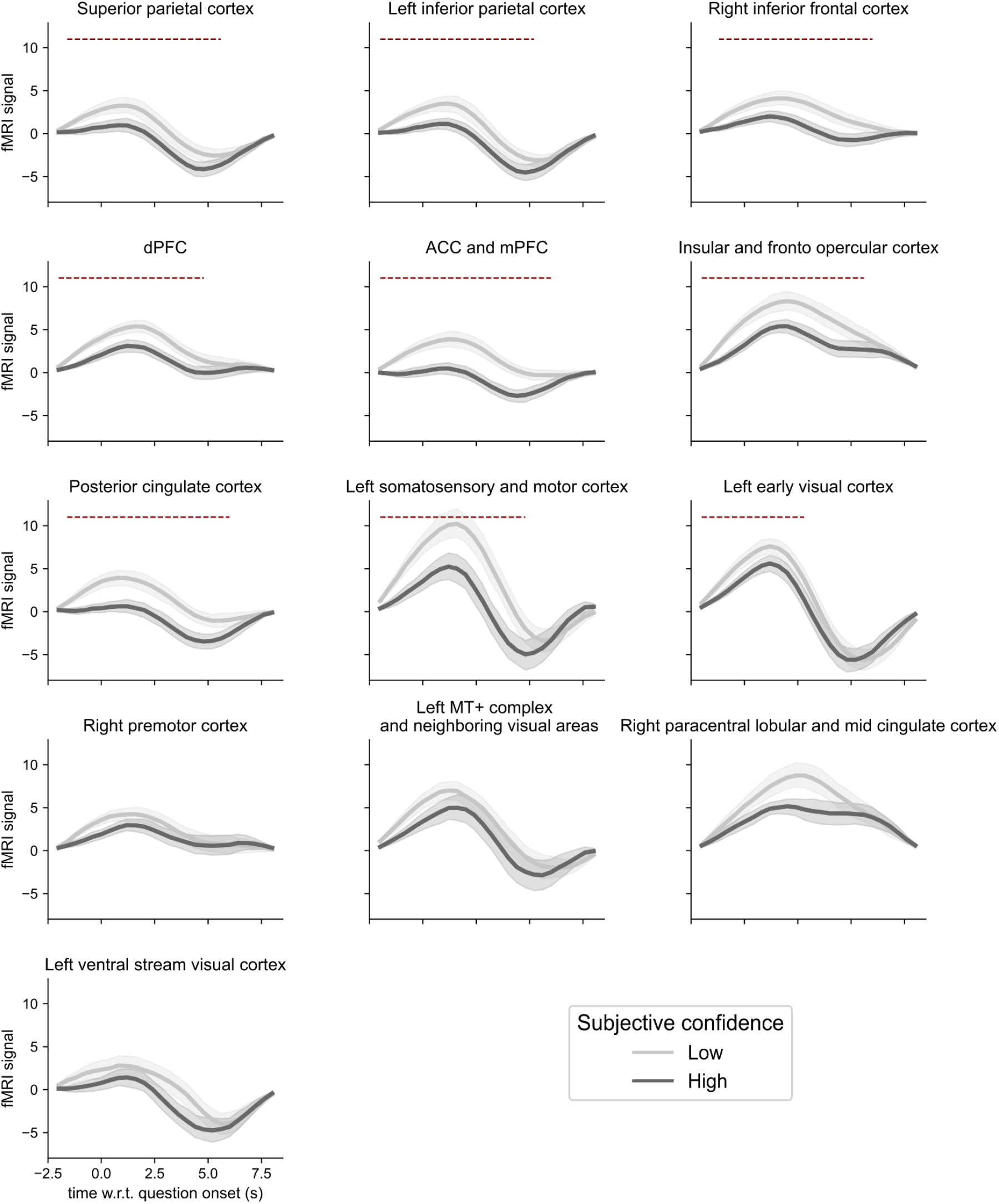
fMRI activity time-course estimated with a FIR model in confidence clusters for low and high subjective confidence. Error shading corresponds to the S.E.M. Red dashed lines indicate clusters with significant difference after multiple-comparison correction with permutations across time (using p<0.05/13 to correct for the 13 ROIs).

## Discussion

Learning in a stochastic and changing environment is difficult but paramount to perception and decision making (Saffran et al., 1996; Behrens et al., 2007; Dehaene et al., 2015; Collins & Shenhav, 2022 for review). Surprise and confidence are suggested to play key roles to dynamically adjust learning in such environments (O’Reilly, Jbabdi, et al., 2013; Faraji et al., 2018; Boldt et al., 2019; Meyniel, 2020; Liakoni et al., 2021). Here, we used a dynamic probability-learning task to investigate the representational level of probabilistic learning in human adults. We identified several regions of frontal and parietal cortices whose activity is sensitive to confidence, in a way that is both specific with respect to the effects of surprise and unpredictability that covaries with confidence, and invariant with respect to which item is predicted. Activity in those regions covaried not only with the optimal level of confidence that should accompany learning, but on top of that with the subjective reports of confidence. Furthermore, in a subset of those regions (the right premotor cortex, the superior parietal cortex, the right inferior frontal sulcus, the intraparietal sulcus, the paracentral lobular and mid cingulate cortex, the right dorsolateral prefrontal cortex, the insular and fronto opercular cortex and the right inferior parietal cortex), the effect of confidence overlapped with the effect of surprise in qualitative agreement with the confidence-weighted principle prescribed by probabilistic inference; those regions could thus be implicated in the human update mechanism at the heart of statistical learning.

### Studying confidence in a learning context

One should keep in mind that confidence is always about something, e.g. ourself (Flavell, 1979; Benabou & Tirole, 2002), a memory (Kelley & Lindsay, 1993; Butterfield & Metcalfe, 2001; Koriat et al., 2002; Carpenter et al., 2019; Li et al., 2021), a decision (Kiani & Shadlen, 2009; Kepecs et al., 2008; Boldt et al., 2019), an estimation (Meyniel, Schlunegger, et al., 2015; Geurts et al., 2022; Tomov et al., 2020). Depending on what confidence is about, it is formalized differently. In studies focusing on confidence about choices (Kepecs et al., 2008; Kiani & Shadlen, 2009; Boldt et al., 2019), *decision* confidence is usually formalized as the probability of the choice being correct (Kepecs & Mainen, 2012; Meyniel, Sigman, et al., 2015; Pouget et al., 2016) and this formalization accounts for subjective confidence (Hangya et al., 2016), although not perfectly (Li & Ma, 2020; Maniscalco et al., 2016; Peters et al., 2017). In contrast, in studies focusing on the confidence about some estimation, such as a learned probability here, the *estimation confidence* intuitively corresponds to the precision of the estimation, and is thus usually formalized as the precision (in the mathematical sense: the inverse of variance) of a posterior distribution (Meyniel, Schlunegger, et al., 2015; Boldt et al., 2019; Yeung & Summerfield, 2012; Navajas et al., 2017). Another formalization could be in terms of the (inverse) entropy of the posterior distribution (Payzan-LeNestour et al., 2013), but it is extremely correlated with precision in our case and thus qualitatively equivalent.

When confidence is formalized as the log precision of a posterior distribution, a model is needed to compute this posterior distribution. Here, we used an ideal observer model that (by definition) inverses the generative process of the task using probabilistic inference to estimate a posterior distribution of the latent item probability based on observations. The optimal confidence associated with this estimate is thus a function of the sequence itself (and the generative process); the origin of a lack of optimal confidence is thus external, not due to internal processes of the observer (Juslin & Olsson, 1997; Walker et al., 2022). This optimal (sequence-related) confidence accounts for the participant’s confidence, but the correlation is far from perfect. Deviations of subjective confidence from optimal confidence could be due to the fact that participants have a different internal model of the generative process, and compute the inference imperfectly or differently (Ma & Jazayeri, 2014; Rahnev & Denison, 2018; Koblinger et al., 2021). Here, we focused on the similarity of optimal and subjective confidence and showed that brain regions sensitive to ideal confidence are also related to the reported confidence, but deviations between the two should be explored in future studies.

### Sensitivity

To study the neural representation of confidence, we made the critical assumption that confidence is represented in some neural activity profile, rather than in synaptic weights (Aitchison et al., 2021). We made this assumption for different reasons. First, measuring synaptic weights requires invasive recording, which is not possible in humans. Second, confidence changes rapidly in the task and can be reported; a neural representation coded in activity levels rather than in synaptic weights could be changed more quickly and read out more easily (Hunt & Hayden, 2017). More specifically, as in most fMRI studies, we assumed a code in terms of activity levels monotonically increasing or decreasing with the encoded variable. This choice formally corresponds to a linear rate code (Dayan et al., 2005) but is mostly made for simplicity and convenience; we do not exclude other forms of coding such as a probabilistic population code (Ma et al., 2006; Ma & Jazayeri, 2014) or a sampling-based code (Hoyer & Hyvärinen, 2003; Paulin, 2005; Fiser et al., 2010; Griffiths et al., 2012). We note that the Probabilistic Population Code predicts under biologically-plausible assumptions that confidence increases with the overall activity level (the “neural gain”) (Ma et al., 2006), which is the opposite of what we found here. Different codes may coexist in different parts of the brain; for instance Geurts et al 2022 found a probabilistic population code for stimulus orientation and the associated confidence in the early visual cortex, and a linear rate code for confidence about the estimated orientation in the frontal lobe.

Other fMRI studies have tested for a representation of confidence with a linear rate code, although they may not use the term “confidence” but instead precision (Iglesias et al., 2013) or the converse notion of uncertainty (Payzan-LeNestour et al., 2013; Tomov et al., 2020; McGuire et al., 2014; Kobayashi & Hsu, 2017). No matter whether those studies involved learning in a context of reward-based decision (Payzan-LeNestour et al., 2013; Tomov et al., 2020; McGuire et al., 2014; Badre et al., 2012), inference (Meyniel & Dehaene, 2017; Kobayashi & Hsu, 2017), perceptual decision (Iglesias et al., 2013), or the estimation of a quantity (e.g. stimulus orientation) without learning (Geurts et al., 2022), they consistently found a negative correlation between confidence and activity in the superior parietal cortex including intraparietal areas, the insular and frontal opercular cortex, the dorsolateral prefrontal cortex (dlPFC), the premotor cortex and the posterior and mid cingulate cortices.

### Specificity

Confidence is often confounded with other quantities. Here we have explored some confounds that relate to the sources of a lack of confidence. One such source is the aleatoric variability of observations (Hora, 1996; Kiureghian & Ditlevsen, 2009). In stochastic environments, observations are generated with intrinsic randomness. This aleatoric variability of observations is quantified as the (un)predictability in our analysis: it is all the stronger that the item probability is extreme. More extreme probabilities lead to more stereotyped sequences (dominated by one item), those probabilities are thus estimated with greater confidence. Confidence correlates with predictability but is not reducible to it because confidence also depends on other factors such as the change point probability that reduces confidence (Behrens et al., 2007). It is important to distinguish confidence from its different contributing factors (e.g. change point probability, which was kept fixed here; unpredictability) because they play different roles in learning (Piray & Daw, 2021; Kalman, 1960; Bland & Schaefer, 2012). The trial-by-trial perceived likelihood of a change point itself (McGuire et al., 2014) is related to surprise (and unexpected uncertainty (Yu & Dayan, 2005)). More surprising observations are more likely to denote the presence of a change point, and confidence decreases after surprising observations indeed. Brain signals that covary with surprise were reported in numerous studies (O’Reilly, Schüffelgen, et al., 2013; Kouider et al., 2015; Mars et al., 2008; Strange et al., 2005). Given the numerous factors that contribute to confidence, or correlate with it, the study of the neural representation of confidence is thus exposed to the issue of specificity (Walker et al., 2022).

### Invariance

We further tested whether the representation of the confidence level that accompanies the latent probability estimate was invariant to the item that was predicted. In the same vein, in another dynamic probability estimation task that presented either images or sounds in different sequences, a representation of confidence in the intraparietal sulcus was found to be invariant to the sensory modality used in the task (Meyniel & Dehaene, 2017). Another study investigated the value (in the sense of pleasure) and the associated confidence that subjects assigned to stimuli from different categories (images of faces, houses and paintings) and found that activity in the ventro-medial prefrontal cortex reflected both the value and confidence in a way that is invariant to the category (Lebreton et al., 2015).

There is an ongoing debate on whether the neural representation of a latent variable and the associated confidence is necessarily a joint representation, or whether the two can be detached from one another in different (and possibly very distant) populations of neurons (Vilares et al., 2012; Walker et al., 2022). In case the representation of confidence is detached, it could be general, invariant to which sensory modality is used, and within a given pair of items, invariant to which item is deemed more likely to occur. In contrast, current formulations of probabilistic population codes (Zemel et al., 1998; Ma et al., 2006; Beck et al., 2008; Ma & Jazayeri, 2014) and sampling-based codes (Hoyer & Hyvärinen, 2003; Fiser et al., 2010; Griffiths et al., 2012; Orbán et al., 2016) assume that the latent variable and the associated confidence are jointly represented in the same activity pattern, which does not favor the invariance property. Testing for invariance is thus informative about the neural code of confidence and the latent variable it accompanies.

### Functionality

One of the strongest tests for functionality is to manipulate the activity pattern putatively subtending a neural representation and assess the changes induced by this manipulation. Weaker tests remain informative about functionality. As highlighted in several works, a key function of confidence in a learning context is belief updating by modulating the effect of surprise (Meyniel, Sigman, et al., 2015; McGuire et al., 2014; Iglesias et al., 2013; Payzan-LeNestour et al., 2013). To identify candidate regions involved in the confidence-weighted update mechanism, we used the fact that if a region were more active when larger updates are made, then it should exhibit greater fMRI signals when surprise is higher and when confidence is lower (O’Reilly, Jbabdi, et al., 2013). We note that the brain regions we identified with this method (the right premotor cortex, the superior parietal cortex, the right inferior frontal sulcus, the intraparietal sulcus, the paracentral lobular and mid cingulate cortex, the right dorsolateral prefrontal cortex, the insular and fronto opercular cortex and the right inferior parietal cortex) were reportedly related to belief updating mechanisms. O’Reilly et al. (O’Reilly, Schüffelgen, et al., 2013) found increased activity in the anterior and posterior parts of the cingulate cortex for larger updates; their analysis did not investigate the effect of confidence on update but demonstrated that the update signal they found in the cingulate cortex is not a mere effect of surprise. This latter study did not find an effect of update in the parietal cortex, but other studies did (Kobayashi & Hsu, 2017). We also found effects of updates in the premotor cortex (comprising the frontal eye field) as in other studies (Vossel et al., 2015; Kobayashi & Hsu, 2017).

The study by Vossel and colleagues makes an interesting link with the concept of attention (Vossel et al., 2015). They used a Posner cueing task with varying cue validity across the experiment and a hierarchical Gaussian filter (HGF) model that included a parameter capturing a precision-dependent attention factor, which reflects in their study the amount of attention allocated to the cued location that modulated belief updating. Selective attention is a top-down modulation of the current sensory input that filters the observations within the sequence, increasing or decreasing their weight. At the mechanistic level, this filtering could correspond to a modulation of the neural gain (Eldar et al., 2013; Marzecová et al., 2018). Selective attention was thus proposed as a mechanism for the regulation of learning (Dayan et al., 2000). Supporting this view, a MEG study (Meyniel, 2020) found a correlation between confidence and beta-band oscillations, which are related to top-down effects (X.-J. Wang, 2010; Siegel et al., 2012) and regulate the extent to which the state of neural networks is maintained or altered by input signals (Spitzer & Haegens, 2017; Engel & Fries, 2010); in line with the confidence-weighting principle, surprise signals were dampened when beta-band power was higher. We also note that the anatomical correlates of surprise and confidence in our study are similar to the ones associated with bottom-up and top-down attention (Corbetta & Shulman, 2002). The causal role of the neural representation of confidence in the regulation of learning and the corresponding mechanisms should be explored in future studies.

## Materials and Methods

### Participants

Twenty-nine (fifteen females) healthy adults with normal or corrected vision, aged between 19 and 37 years old (mean 25.41, SEM 1.02), participated in the experiment. The study was approved by the local ethics committee (references: CPP-100032 and CPP-100055 Ile-de-France) and all participants gave written informed consent prior to participating. Data from three subjects were discarded because of acquisition problems, resulting in a final sample size of 26 participants.

### Stimuli

The set of stimuli consisted of Gabor patches with an orientation of −45 or +45 degrees. The phase of the Gabor patch was randomly drawn on each trial. All stimuli were perceived and distinguished without ambiguity. They were presented on a screen at 200 cm from the participants (visual angle=2.60 x 2.60 degrees; frequency of the grating=3.2 cycles per degree; Gaussian standard deviation=0.42 degrees).

### Experimental design

Participants performed a probability-learning task (see Figure 1, upper section). In each session, they were shown a binary sequence of 420 stimuli, each displayed for 1 s with an inter-stimulus interval of 300 ms that presented a fixation cross. The two possible items in the sequence were two Gabor patches with different orientations, which we denote A and B here for simplicity (see “Stimuli”). Sequences were generated based on a Bernoulli process whose parameter (the item probability θ = *p(A))* changed abruptly, unpredictably, and without notice. Subjects were asked to covertly estimate the item probability and its changes at all times, and to report occasionally their probability estimate and the associated confidence when prompted by a question screen (every 22 stimuli on average with a jitter of ± 1, 2 or 3 stimuli). Those questions aimed to assess probability learning in subjects and to promote engagement in the task.

Subjects first performed one training session (not included in the analysis) outside of the scanner where they reported probabilities and confidence on a quasi-continuous scale by moving a cursor. Then they performed four sessions in an MRI scanner with a simplified reporting procedure: the reporting scale was discretized into bins, each corresponding to a dedicated key press on a response box. This reporting procedure limited response times (which were let free) and motor actions (since a single key press suffices for each scale) during fMRI recording. Despite the discretization of the scales, subjects were verbally encouraged to estimate a continuous probability estimate as they did during training. Half of the subjects reported their probability estimate and then their confidence on two five-bin scales. For the other half of the subjects, the order of the questions was reversed (confidence, then probability estimate) and the probability scale used either 3 or 5 bins randomly across trials in order to prevent participants from preparing a motor response during their covert probability estimation.

The item probability remained constant between change points in a sequence. Change points could occur independently of one another on any given stimulus with a probability 1/75. The number of stimuli in a stable period thus follows a geometric distribution, with an average of 75 stimuli; to avoid exceedingly long stable periods, we truncated this distribution to a maximum of 300 stimuli. On the first stimulus, the item probability was sampled uniformly in the interval [0.1, 0.9], and when a change point occurred it was resampled in the same interval with the constraint that the change in odd ratios should be at least fourfold, i.e. that 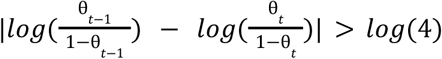. Before the experiment, subjects were fully informed in a non-technical way about the generative process of sequences, except for the numeric values of the change point probability (which was just said to be “rare”), and the constraints on the sampling of item probabilities.

The task was programmed and run using Python (version 3.7) and Expyriment (0.10.0).

### The Ideal Observer: An optimal Bayesian model of learning

The ideal observer is a model of learning that provides optimal observation-driven estimates of the changing item probability θ*_t_*. More precisely, the ideal observer estimates the likelihood of current item probability *p*(θ*_t_*|*y*_1:*t*_) given past observations (*y*_1:*t*_) using probability calculus, and in particular, Bayes’ rule. In the case of an abruptly changing Bernoulli process, this ideal observer is well known (Meyniel et al., 2016; Gallistel et al., 2014). The ideal observer starts from a flat prior on the item probability θ_0_ and leverages the Markov property of the generative process to update its estimate after each new observation: the next item probability θ_*t*+1_ only depends on its current value θ*_t_* (they are equal if no change point occurs, and different otherwise). This property, together with the fact that the likelihood of the current stimulus only depends on the current item probability, make it possible to update the item probability by moving forward in the sequence:

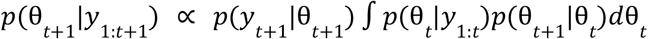

Note that *p*(*y*_*t*+1_|θ_*t*+1_) = θ_*t*+1_ if *y*_*t*+1_ = *A* and 1 – θ_*t*+1_ otherwise, by definition of a Bernoulli process. The transition probability of the item probability is captured by *p*(θ_*t*+1_|θ*_t_*); our ideal observer knows the true probability of change point (1/75) and considers that the item probability is sampled uniformly in the range [0, 1] when a change point occurs (it is thus unaware of the other constraints of the generative process, like participants).

Given the estimation of the item probability, the prediction for the next stimulus is:

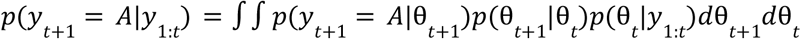

We formalized the confidence about this prediction as the log-precision of the posterior distribution (Meyniel, Schlunegger, et al., 2015; Boldt et al., 2019; Meyniel, Sigman, et al., 2015; Yeung & Summerfield, 2012; Navajas et al., 2017):

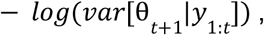

Following Shannon (Shannon, 1948) we formalized surprise as the log-improbability of the item observed on trial *t* + 1:

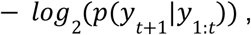

and the unpredictability as the entropy of the prediction:

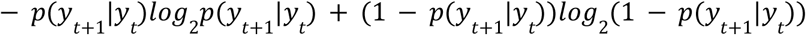

The ideal observer used the sequences of stimuli shown to the subjects. The probability estimates and confidence of participants and the ideal observer were compared at the moment of questions.

### MRI data collection and preprocessing

MRI data were acquired on a Siemens MAGNETOM 7 Tesla MRI scanner (Siemens Healthcare, Erlangen, Germany) with whole-body gradient and 32-channel head coil (Nova Medical, Wilmington, MA, USA). Functional MRI data were T2*-weighted fat-saturation echo-planar image (EPI) volumes with 1.5 mm isotropic voxels, recorded using a multi-band sequence (Moeller et al., 2010) (https://www.cmrr.umn.edu/multiband/) with the following parameters: multi-band factor [MB] = 2, GRAPPA acceleration factor [IPAT] = 2, partial Fourier [PF] = 7/8, matrix =130 x 130, number of slices = 68, slice thickness =1.5 mm, repetition time [TR] = 2 s, echo time [TE] = 22 ms, echo spacing [ES] = 0.71 ms, flip angle [FA] = 68°, bandwidth [BW] = 1832 Hz/px, phase-encoding direction from anterior to posterior. The calibration preparation was done using Gradient Recalled Echo (GRE) data. At the beginning of each scanning experiment, a B0 map was acquired and loaded in the console for shimming, first of the whole brain, then in an interactive fashion on the occipito-parietal cortex, aiming to reduce the FWHM and increase the T2*. Subsequently, a B1 field map was acquired, on the basis of which the system voltage was chosen according to the reference voltage computed from the B1 map values in the intraparietal sulcus region. Before each session, two single volumes were acquired with the parameters listed above but with opposite phase encode directions in order to be used later for distortion correction (see “MRI data analysis”). T1-weighted anatomical images were acquired with 0.75 mm isotropic resolution using an MP2RAGE sequence, with the following parameters: GRAPPA acceleration with [IPAT] = 3, partial Fourier [PF] = 6/8, matrix = 281×300, repetition time [TR] = 6 s, echo time [TE] = 2.96 ms, time of inversion [TI] 1/2 = 800/2700 ms, flip angle [FA] 1/2 = 4°/5°, bandwidth [BW] = 240 Hz/px. Head movement was minimized by padding and tape. Visual stimuli were displayed on the MRI-compatible LCD screen (BOLDscreen 32 LCD for fMRI, Cambridge Research Systems; resolution=1920×1080; refresh rate=120 Hz) at the back-end part of the scanner (viewing distance=200 cm) and viewed through a mirror fixed onto the head coil. Participants reported responses using their right hand and a fiber optic response pad (fORP, Current Designs, Philadelphia, PA, USA) featuring five buttons.

EPI images were corrected for slice-timing (with respect to the middle temporal slice of a volume), for motion with rigid transformations, and co-registered to the anatomical T1-weighted MP2RAGE image using the Statistical Parametric Mapping software (SPM12 https://www.fil.ion.ucl.ac.uk/spm/software/spm12/). The single-band reference images of the two volumes acquired prior to each session with opposite phase encoding directions were used to estimate a set of field coefficients (FSL TOPUP https://fsl.fmrib.ox.ac.uk/fsl/fslwiki/FSL/), which was then used to apply distortion correction (FSL apply_topup) to EPI images of the corresponding session. The preprocessing steps relying on FSL and SPM used the Python interfaces provided by Nipype (https://doi.org/10.5281/zenodo.596855). Time series in each voxel were detrended, high-pass filtered (at 1/128 Hz) and z-scored across experimental runs.

For the purposes of surface-based analyses, each subject’s cortical surface was segmented using Freesurfer (https://surfer.nmr.mgh.harvard.edu/) based on the anatomical MP2RAGE image, and normalized to the fsaverage7 template (163,842 vertices per hemisphere). The same transformation was used to process the functional data: for all analyses, coefficient maps were first estimated in each single subject’s volume space, then projected onto the native reconstructed surface with Freesurfer, then aligned to the high-resolution fsaverage7 mesh and finally smoothed with a 3-mm FWHM Gaussian kernel.

### MRI data analysis

Statistical analyses of MRI data were performed using general linear models (GLM) that included regressors of interest along with realignment parameters as covariates of no interest.

In GLM_1_, we modeled the stimuli onsets as an indicator variable and several parametric modulations of it as separate regressors (the prediction about what the current stimulus would be, the corresponding unpredictability and confidence, and the surprise arising from the stimulus actually presented, all derived from the ideal observer), the question onsets as an indicator variable and several associated parametric modulations of it as separate regressors (the ideal observer’s prediction and confidence as well as the subject’s prediction and confidence in response to the question) and realignment parameters.

GLM_2_ was similar to GLM_1_, with the difference that we included separate regressors for confidence signals depending on whether the prediction favored stimulus A or stimulus B (p≤0.5 or p≥0.5 respectively).

GLM_1_ and GLM_2_ were estimated in the native space (non-normalized volumes) of each subject, and the fitted regression coefficients in each voxel were then projected onto the cortical surface and normalized.

GLM_3_ was similar to GLM_2_ with the difference that the separate regressors for confidence signals were also split according to the confidence values in 6 different bins.

GLM_4_ corresponds to the Finite Impulse Response (FIR) analysis of subjective confidence around the question onset. The input fMRI time-series were extracted from the clusters exhibiting an effect of the ideal confidence (identified with GLM_1_, those clusters are displayed on Fig 2A). The fMRI signal was averaged across vertices for each cluster and for each subject and was temporally upsampled by linear interpolation (factor 5). For each subject, questions were sorted into those having low vs. high confidence reports (using a subject-specific threshold that minimized the sample size difference between those two categories) and FIR regressors were generated for each category within a time window of −2 s to 8 s relative to the question onset. Note that each question appears only 1300 ms after the onset of the previous stimulus and that confidence reports correlate with the ideal observer’s confidence, the question-locked FIR estimates could thus capture an effect of the ideal observer’s confidence (or another quantity) during stimulus presentation. In order to avoid this potential confound, GLM_4_ also included convolved regressors corresponding to the stimulus onset and its parametric modulation by the ideal observer’s prediction, unpredictability, confidence, and surprise as in GLM_1_. The goal is to test whether fMRI activity around the question onset reflects subjective confidence, above and beyond the effect of ideal confidence that is present during the sequence presentation.

For the sake of simplicity, neighboring and homologous clusters of confidence and overlap between confidence and surprise were merged for GLM_2_, GLM_3_ and GLM_4_. The grouping was based on criteria defined in the HCPMMP1.0 atlas’ neuroanatomical results (Glasser et al., 2016, Supplementary Neuroanatomical Results For A Multi-modal Parcellation of Human Cerebral Cortex).

### Simulation: parameter recovery analysis

We simulated 1000 experiments (each consisting of four sessions of 420 observations) with the same timings as participants and used the ideal observer model to construct GLM_1_ (omitting the questions-related and motion-related regressors since no participant data are available in simulations). Then, fMRI-like activity was simulated as a weighted linear sum of regressors (sampling the weights independently and uniformly in the interval [0-1]) and then corrupted with autocorrelated Gaussian noise (the SD was adjusted so that the ratio between the noise-free and noisy signals was 0.10 on average, and we introduced a noise autocorrelation of 0.90 using an AR(1) model, which is a very pessimistic case since the median autocorrelation (across voxels and subjects) found in our fMRI data in the gray matter was 0.16. GLM_1_ was then fitted onto the simulated noisy fMRI BOLD activity. To quantify the recovery, we computed Pearson’s correlation between the generative and fitted weights assigned to each regressor.

### Overlap analysis

ROIs were defined based on the group-level overlap of the clusters corresponding to surprise and uncertainty (negative effects of confidence) in GLM_1_ on the surface (fsaverage). For each individual and each ROI, the regression coefficients associated with surprise and uncertainty were extracted for each vertex. We then computed the Pearson’s correlation (ρ) between the regression coefficients of surprise and uncertainty across vertices, in each ROI and each participant. We then z-scored the observed Pearson’s correlation (z-ρ) using the mean and SD of a null distribution of ρ values. We simulated a null distribution (50 samples) for each participant and ROI by repeating the estimation of ρ as above, but after substituting the sequences of stimuli observed by the participant with another set of sequences (randomly selected).

### Statistical analysis

Individual surface-based, normalized maps of regression coefficients entered a group level analysis: a t-test against 0. The resulting statistical maps were thresholded at a cluster-forming threshold of p < 0.001 and corrected for multiple comparisons across vertices at the cluster level at p_FWE_ < 0.05 (Monte Carlo-based cluster correction from FreeSurfer).

FIR estimates in each cluster entered a paired t-test testing for the difference between low vs. high confidence reports. Peri-question time points exhibiting a difference (two-sided test) significant at p<0.05 formed clusters, and cluster-level p_FWE_ (correcting for multiple comparisons across time points) were estimated using 10,000 permutations (permutation_cluster_1samp_test from MNE Python (Gramfort, 2013)). Additionally, these results were corrected for multiple comparisons across the ROIs with a Bonferroni correction, by dividing the threshold p=0.05 by the number of ROIs.

For the overlap analysis, the subject-level z-ρ values were tested for significance at the group level with a t-test against 0. The results were corrected for multiple comparisons across the ROIs with a Bonferroni correction.

## Acknowledgements

We thank all recruited volunteers who participated in the study, the fMRI staff of the NeuroSpin center (Véronique Joly-Testault, Laurence Laurier, Gaëlle Médiouni, Valérie Berland, Chantal Ginisty, Yann Lecomte), Isabelle Denghien and Christophe Pallier for advice in the fMRI data preprocessing.

This work was funded by the French National Research Agency (grant ANR #18-CE37-0010-01 CONFI-LEARN to F. Meyniel), the European Research Council (grant ERC #947105 StG NEURAL-PROB to F. Meyniel) INSERM, the Leducq Foundation (ERPT Equipment Program for the 7T MRI clinical device), the Agence de l’Innovation pour la Défense (grant awarded to T. Bounmy) and the Ecole Doctorale FIRE – Programme Bettencourt.

## Additional information

### Authors contributions

T.B.: Experiment code, data acquisition, MRI data preprocessing and analyses, interpretation, writing; E.E.: fMRI acquisition procedures and data analyses (validation); writing (review); F.M.: Conceptualization, supervision, validation, fMRI data analyses, writing (review and editing).

### Ethics

The study was approved by the local ethics committee under the references CPP-100032 Ile-de-France VII and CPP-100055 Sud-Ouest et Outre Mer III. All participants gave their informed written consent before participating in the study.

### Code availability

The Python code of the (ideal) Bayesian observer can be found at https://github.com/florentmeyniel/TransitionProbModel/.

## Supplementary Materials

**Figure 1—figure supplement 1.**
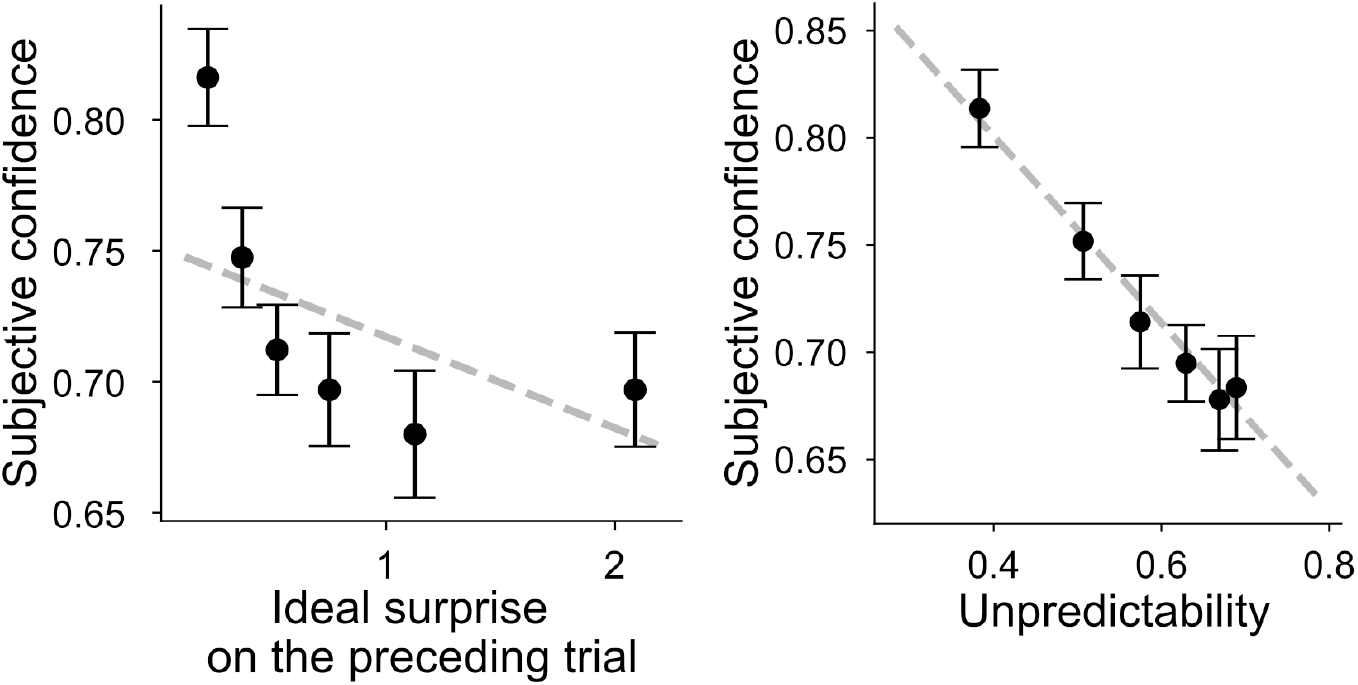
Subjective levels of confidence exhibit different qualitative signatures of the Bayesian solution. Subjective confidence plotted against the ideal observer surprise on the preceding trial (left) or unpredictability (right). The dashed lines correspond to the mean linear fits obtained at the subject level. Bins correspond to 6 percentiles of surprise on the preceding trial or unpredictability.

**Figure 2—figure supplement 1.**
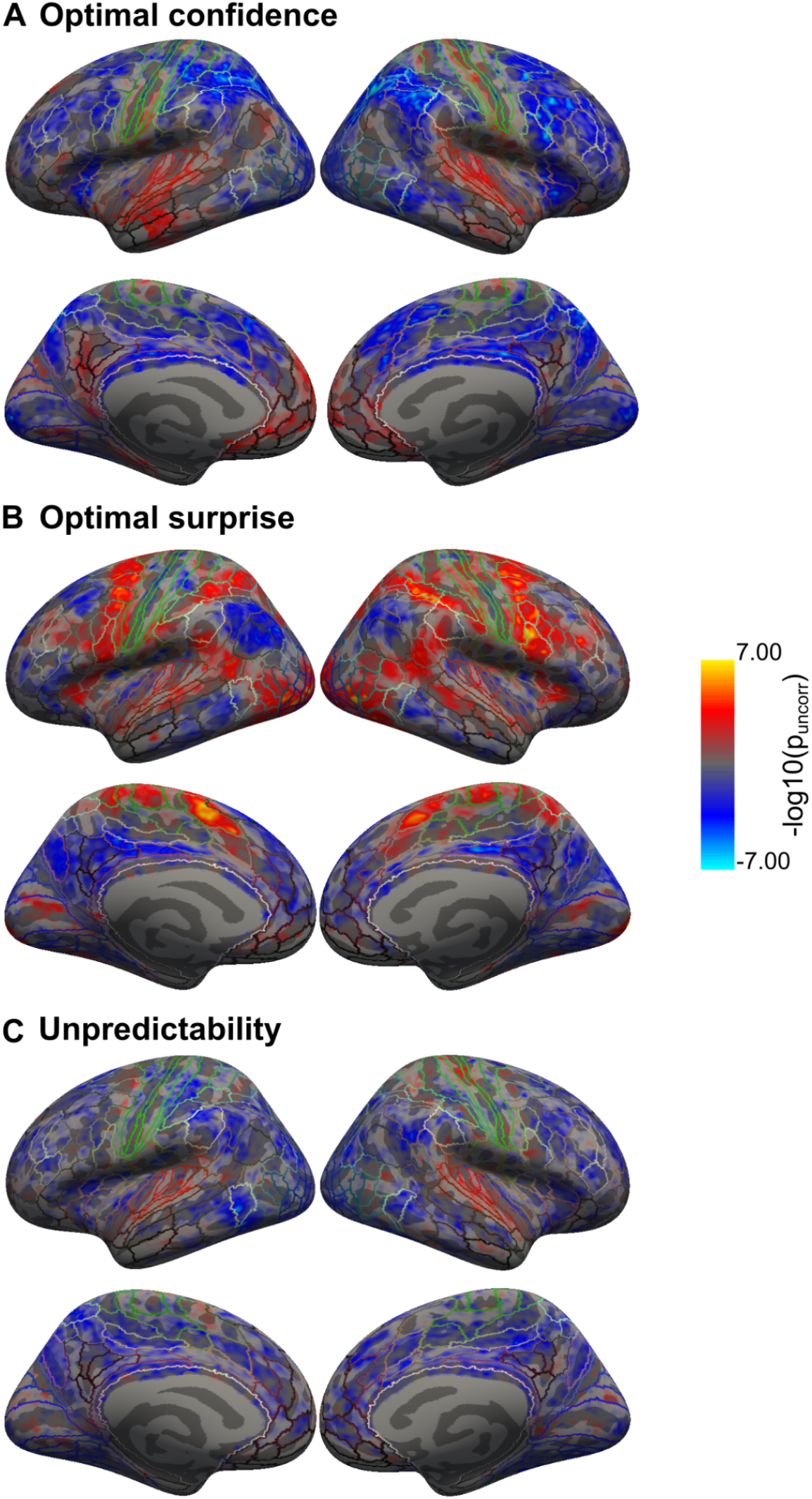
Group level significance maps for confidence, surprise and unpredictability. Cortical brain maps are displayed on Freesurfer’s fsaverage surface with delineations based on the HCP-MMP1.0 atlas (Glasser et al., 2016).

**Figure 2—figure supplement 2.**
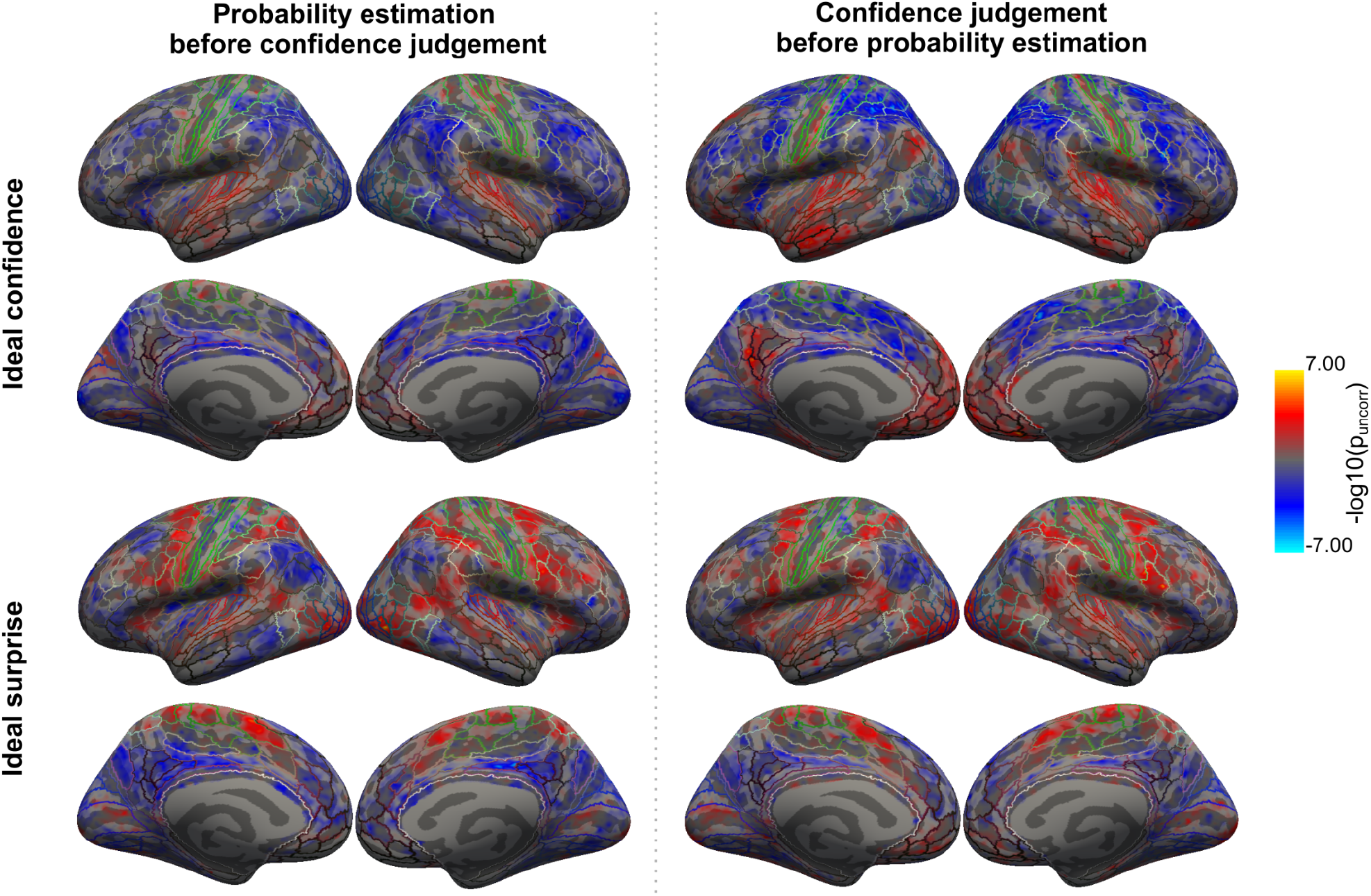
Identified cortical networks of confidence and surprise in the two groups of subjects with different behavioral reporting structures. Cortical correlates of confidence and surprise identified in the subgroup that first estimated probabilities then reported their prediction confidence both on a five-step scale (left) are similar to the networks identified in the subgroup that first provided prediction confidence on a five-step scale, then reported the probability estimate on a three- or five-steps scale (right). Cortical brain maps are displayed on Freesurfer’s fsaverage surface with delineations based on the HCP-MMP1.0 atlas (Glasser et al., 2016).

**Figure 2—figure supplement 3.**
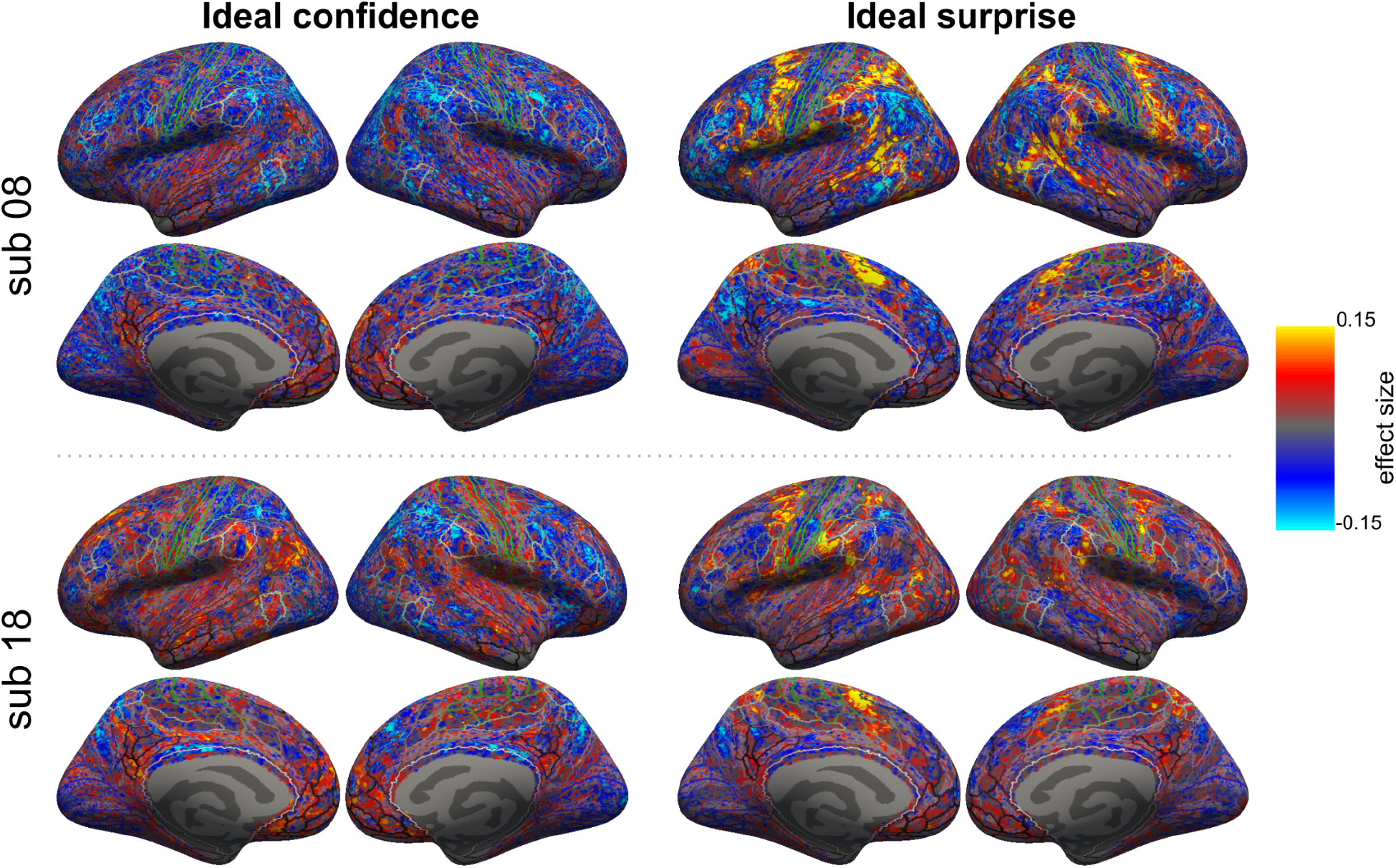
Example subjects. Subject level coefficient maps of confidence and surprise.

**Figure 3—figure supplement 1.**
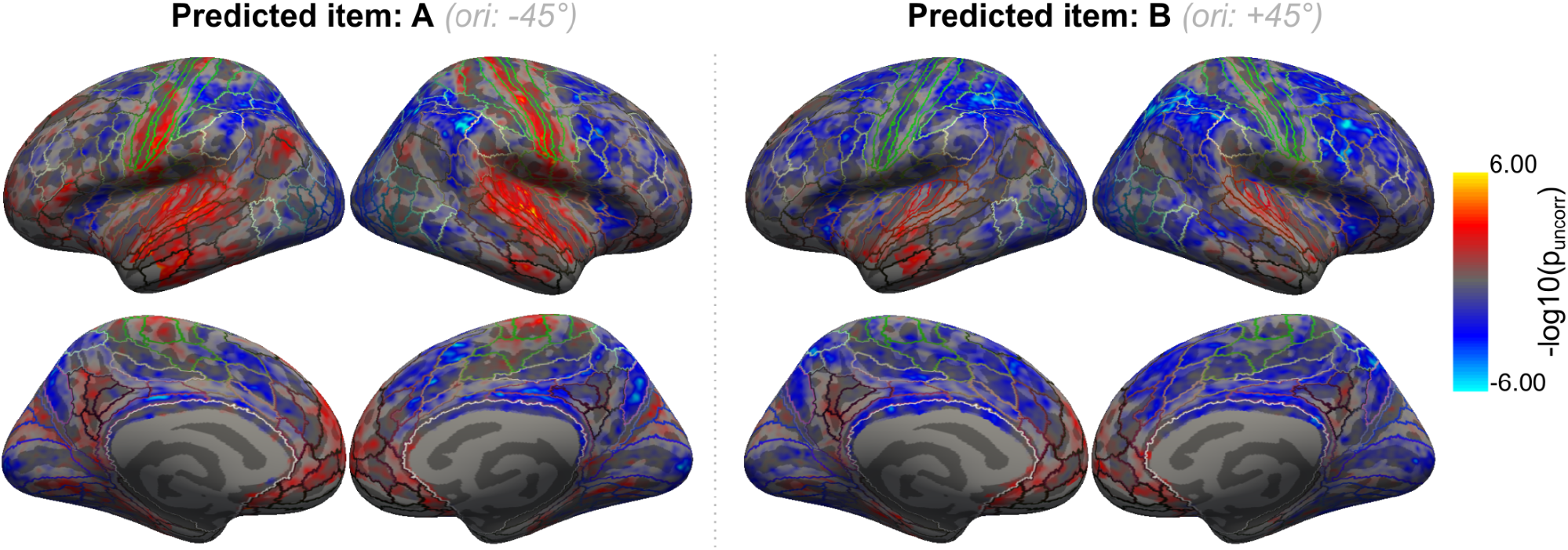
Neural correlates of confidence associated with the prediction of a Gabor patch oriented counterclockwise (stimulus A, orientation: −45°) or clockwise (stimulus B, orientation: +45°). Cortical brain maps are displayed on Freesurfer’s fsaverage surface with delineations based on the HCP-MMP1.0 atlas (Glasser et al., 2016).

